# An integrative spatial multi-omic workflow for unified analysis of tumor tissue

**DOI:** 10.1101/2024.10.15.618574

**Authors:** Jurgen Kriel, Joel J.D. Moffet, Tianyao Lu, Oluwaseun E. Fatunla, Vinod K. Narayana, Adam Valkovic, Ana Maluenda, Malcolm J. McConville, Ellen Tsui, James R. Whittle, Sarah A. Best, Saskia Freytag

## Abstract

Combining molecular profiling with imaging techniques has advanced the field of spatial biology, offering new insights into complex biological processes. Focusing on diffuse *IDH*-mutated low-grade glioma, this study presents a workflow for Spatial Multi-omics Integration, SMINT, specifically combining spatial transcriptomics and spatial metabolomics. Our workflow incorporates both existing and custom-developed computational tools to enable cell segmentation and registration of spatial coordinates from both modalities to a common coordinate framework. During our investigation of cell segmentation strategies, we found that nuclei-only segmentation, while containing only 40% of segmented cell transcripts, enables accurate cell type annotation, but does not account for multinucleated cells. Our integrative workflow including cell-morphology segmentation identified distinct cellular neighborhoods at the infiltrating edge of gliomas, which were enriched in multinucleated and oligodendrocyte-lineage tumor cells, that may drive tumor invasion into the normal cortical layers of the brain.

**Highlights:**

- Alignment and integrated analysis of spatial transcriptomic and metabolomic data
- Nuclei-only and cell-morphology segmentations are concordant for cell annotation
- Spatially distinct regions are conserved in transcriptomic and metabolomic datasets
- Multi-omic exploration of glioma leading edge identifies novel biological features

## Introduction

Spatial biology unlocks tissue complexity by providing the context in which cells and molecules exist. Imaging-based methods such as spatial transcriptomics (ST) and spatial metabolomics (SM) enable the mapping of molecular features to spatial locations. Using these technologies, researchers have gained a more comprehensive understanding of biological processes such as embryonic development, tissue morphogenesis and tumorigenesis^1–5^. Despite providing spatial context, the application of a single technology is unlikely to uncover regulatory mechanisms occurring across several molecular levels. Integrating data generated from multiple technologies has the potential to reveal a more accurate model of the complex interplay of molecular regulators. This integration is particularly valuable in the study of complex diseases, such as glioma, where heterogeneous non-uniform cells reside in a complex microenvironment, thus understanding regulatory pathways within their spatial context is critical for understanding the disease and identifying potential therapeutic targets. However, two challenges – cell segmentation and alignment - hinder the effective combination of spatial data from multiple technologies and their ability to inform on cellular context.

Cell segmentation is a foundational step to assign transcripts and extract meaningful cellular information from ST data but lacks optimal solutions. The accuracy of cell segmentation can vary significantly depending on tissue type, ST platform, segmentation approach, and data compression strategies^1,2,6,7^. Cell segmentation accuracy is most impacted by how cell boundaries are determined. This ranges from nuclei expansion strategies, which estimates cell boundaries based on nuclear size and location (10x Genomics Xenium versions 1.1-1.9), to thresholding of cell outlines based on fluorescence staining to reflect true cell morphology more accurately (NanoString CosMx, Vizgen MERSCOPE, 10x Genomics Xenium versions 2-3.1.0 additional kit). Despite advancements, proprietary staining methods are not universally optimized for all tissue types, complicating standardization of cell segmentation across studies. This challenge is exacerbated in tissues with morphologically diverse cell types, such as glioma, necessitating the use of sophisticated segmentation models. For example, while immune cells, due to their roughly spherical cell shape, are appropriately estimated by expansion-based segmentations, branching cell types including neurons and astrocytes are not^8–10^. Furthermore, the presence of multinucleated cells is ignored by many segmentation methods, including nuclei expansion segmentation, which all assume each nucleus exists as a distinct cellular entity.

Next, the alignment of multi-modal datasets requires the establishment of a common coordinate framework, where the initial coordinates of each modality are transformed such that there is spatial correspondence between them. This process can be challenging due to non-linear distortions, varying scale and orientation, and non-uniformity across sections^11^. Furthermore, when investigating associations between metabolites and transcripts on serial sections, this alignment must be performed with micrometer precision to enable resolution at the cellular level. While alignment strategies commonly leverage landmark features of tissues, recent advancements have enabled the use of iterative gradient descent to improve registration^12^. It is critical that workflows ensure proper alignment of tissue to avoid identifying improper associations across multi-modal data.

Here, we address these challenges and present Spatial Multi-omics Integration (SMINT), a unified analysis framework for the effective analysis of ST (10x Genomics Xenium, 339 gene custom panel) and SM (MALDI-IMS)^13^ data generated from sequential sections of surgically resected low-grade glioma. The morphological diversity of glioma, which contains spherical and branching cell types, offers a representative tissue type that requires addressing one of the overarching challenges associated with segmentation and alignment of any tissue profiled with multi-omics spatial technologies. This allows us to vigorously test our framework and investigate segmentation performance by assessment of cell type characteristics captured – including morphology, annotation, and multinucleated status. Furthermore, mutation in the enzyme isocitrate dehydrogenase (IDH), ubiquitous in low-grade glioma, results in the production of oncometabolite 2-hydroxyglutaric acid (2-HG)^14^. Overall, the study of glioma architecture can benefit from the combination of ST and SM. We demonstrate that the SMINT workflow for integrative analysis across technologies – combining transcriptomic profiles, metabolic signatures, tissue architecture, and immunofluorescence – reveals biological pathways governing glioma with multi-level validation.

## Results

### Post staining allows for cell-morphology segmentation

To examine spatial modalities in low-grade glioma, we sectioned a WHO CNS grade 3 *IDH-*mutant astrocytoma. Neuropathology review identified features of central tumor transitioning to healthy brain tissue, with a characteristic infiltrating edge at the intersection (**Figure 1a, Figure S1a**). Serial sections (10 μm thick) were generated for hematoxylin and eosin (H&E) stain, 10x Genomics Xenium on fiduciary slides and MALDI-IMS on indium tin oxide (ITO) coated glass slides. ST was performed using the 10x Genomics Xenium platform with 339 custom probes (**Table S1**), followed by post-staining immunofluorescence (IF) with glial fibrillary protein (GFAP), Phalloidin, and DAPI to visualize cell membrane boundaries (**Figure 1b-c**). This setup allowed us to explore differences and similarities between three different segmentation strategies: nuclei-only, expansion-based, and cell-morphology segmentation using Cellpose^6^ (**Figure 1d-g**, **Figure S1b-g**). Both nuclei-only and cell-morphology captured less cells than expansion-based segmentation, though this was partially due to regional tissue distortions in the post-staining process (Figure **S1h**). The expansion-based segmentation method showed poor correspondence to the cellular architecture revealed by IF, whereas the custom cell-morphology segmentation displayed refined demarcation of cell boundaries (**Figure 1d-g**). Morphometric analysis indicated that the cell area of expansion-based segmentation was on average 3 times larger than matched cell-morphology segmentation (**Figure 1h**). For expansion-based segmentation, cell area was strongly correlated with the number of neighboring cells within 15 µm of the cell centroid (−0.76), whereas there was weak correlation for cell-morphology segmentation (- 0.15). While both segmentation strategies produced cells of comparable roundness, this value was more correlated with neighboring cell number for the expansion-based method (Expansion-based: moderate correlation, −0.44. Cell-morphology: weak correlation, −0.16) (**Figure 1i**). These results highlight that expansion-based segmentation leads to overly expansive cell boundaries and implausible morphology that are primarily caused by local cell density. In contrast, the cell-morphology model produced more reliable cell masks that account for cellular geometry.

**Figure 1.**
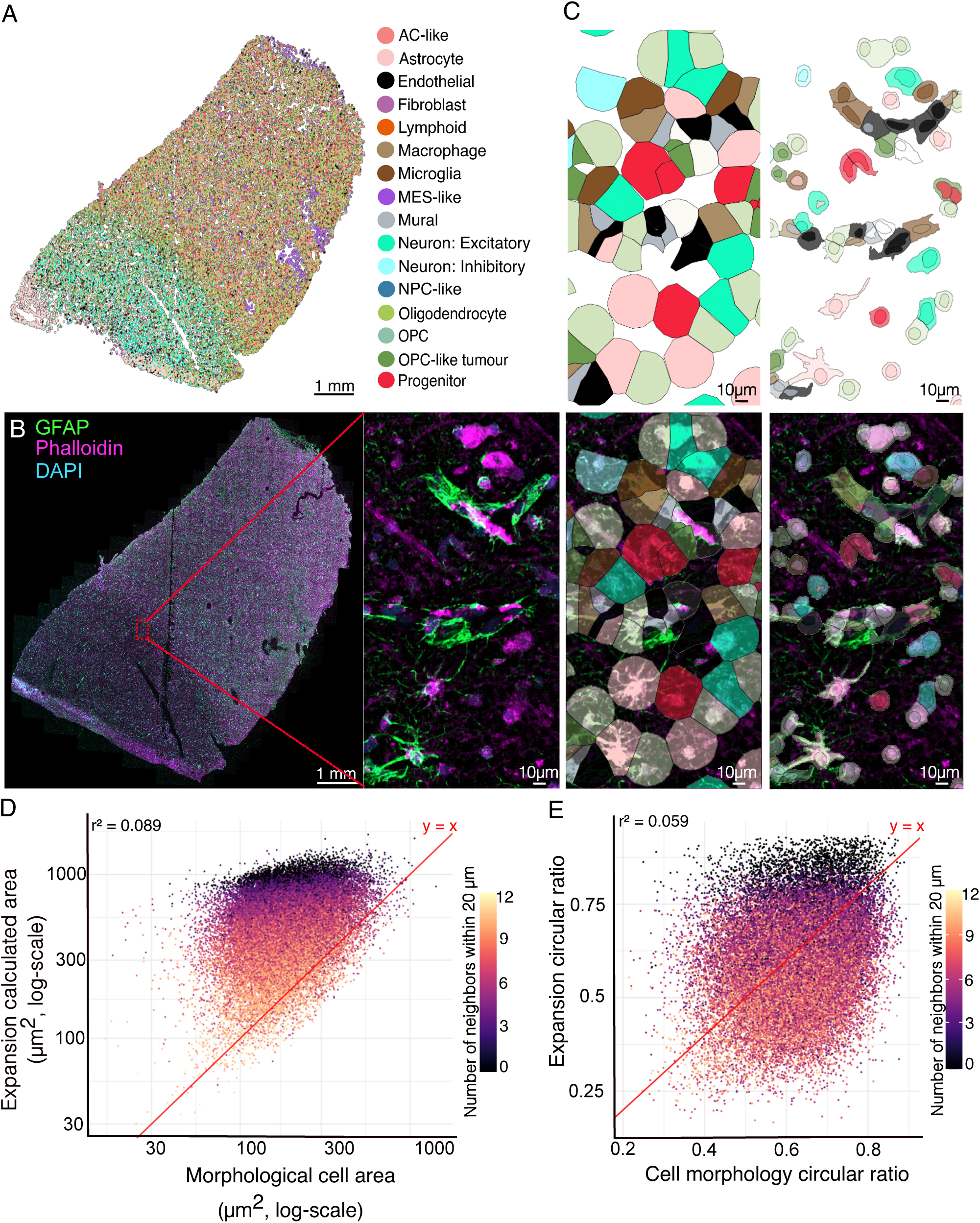
Cell-morphology segmentation identifies realistic cell shapes. **A.** Distribution of cell types annotated by nuclei-only and cell-morphology segmentation. Sample annotated by cell type, showing only nuclei which share the same cell type annotations with cell-morphology segmentation. **B.** IF tile scan conducted post 10x Genomics Xenium data acquisition, showing cells stained for glial fibrillary protein (GFAP, green), actin (Phalloidin, magenta) and nuclei (DAPI, cyan). Scale, 1 mm. Inset indicated in red box, used in C. Scale, 50 µm. **C.** Structural comparison between polygons generated from expansion-based segmentation (top, left), overlaid with IF (bottom, left) and nuclei-only with overlaid cell-morphology segmentation (top, right), overlaid with IF (bottom, right). Scale, 10 µm. **D.** Comparison of cell area between expansion-based and cell-morphology segmentation strategies, only considering cells identified by both segmentations. The x-axis is the size of cell area as estimated by cell-morphology and the y-axis is the size of cell area estimated by expansion-based segmentation. Points are colored by the average number of nearest neighbors within 15 µm between the two segmentation strategies. **E.** Comparison of circular ratio between expansion-based and cell-morphology segmentation strategies for matched cells identified by both segmentations. This ratio for a particular cell mask is calculated by dividing the area of the maximum inscribed circle by the area of minimum binding circle. The x-axis is the ratio based on cell-morphology segmentations, and the y-axis is the ratio based on expansion-based segmentations. Cells are colored by the average number of nearest neighbors within 15 µm between the two segmentation strategies.

### Segmentation strategies result in highly concordant cell type annotation

The choice of segmentation strategy dictates the assignment of transcripts, thereby affecting cells’ transcriptomic profile and subsequent cell type annotations. Hence, we explored the effect of the segmentation strategies on cell annotation as a readout of transcript assignment efficacy. These annotations were performed by semi-supervised clustering, cell type prediction and marker expression using multiple references of single cell glioma populations^15,16^. Paired overlapping cells from the expansion-based and cell-morphology segmentations were assessed for concordance of cell type annotations. Although cell-morphology and expansion-based segmentation differ in estimated cell size, there was 88% concordance between their annotations. Next, we assessed nuclei-only segmentation compared to cell-morphology segmentation. Remarkably, despite containing just 40% of the segmented cells’ transcripts, annotations based on the two approaches were 87% concordant (**Figure 2a, Figure S2a-g**). Discordant annotations could be attributed to three key features. One quarter were between cell types with highly correlated transcript signatures (>0.75), indicating closely related cell states (**Figure 2a, Figure S2h**). Discordant annotations were associated with a lower proportion of transcripts bound within the nucleus, reaffirming that larger differences in transcriptomic profile between nucleic and cytoplasmic spaces cause less concordance (**Figure S2i**). In regions that would typically be excluded from further analysis, including where the tissue had folded-over during processing (**Figure 2b-c, Figure S1a**), discordant annotation was due to significant overlap of cells. Discordant annotations represented 25% of nuclei in this folded region. In general, nuclei-only segmentation resulted in the overrepresentation of mesenchymal-like tumor cells and excitatory neurons compared to the cell-morphology annotation, likely due to increased transcript contained in the nucleus; whereas in cell-morphology segmentation oligodendrocytes, microglia and vasculature were enriched, likely due to their non-circular morphology (**Figure S2j**).

**Figure 2.**
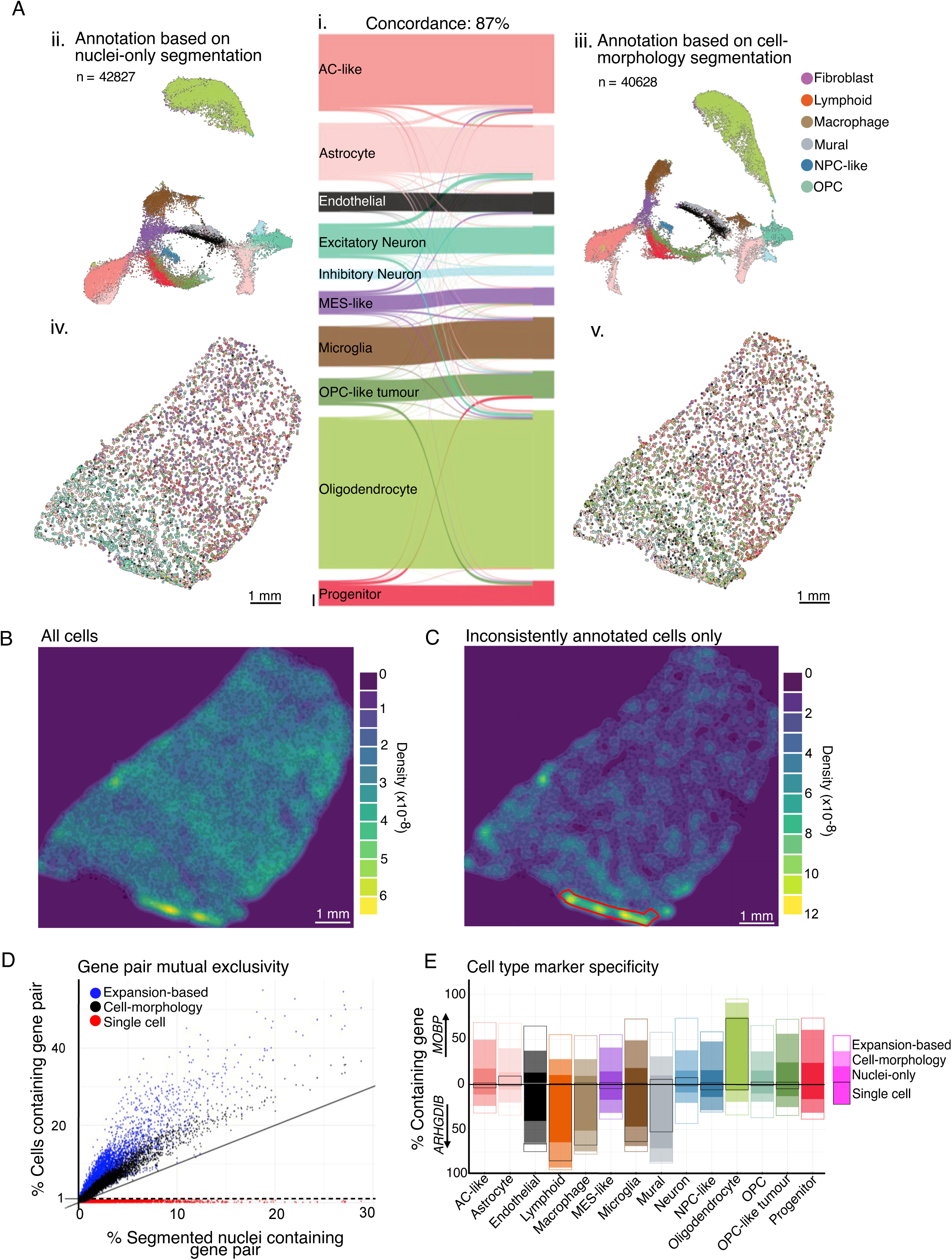
Concordance of cell type annotations using nuclei-only and cell-morphology segmentation strategies. **A.** i) Sankey plot showing change in annotation for matched segmented nuclei (left) and cells (cell-morphology segmentation, right) of highly abundant cell types. Scale bar on bottom left, 1000 segmentations. ii) UMAP of nuclei-only segmented cells, colored by cell type annotation. iii) UMAP of cell-morphology segmented cells, colored by cell type annotation. iv) Distribution of inconsistently annotated cells, colored by cell type annotation from nuclei-only segmentation. v) Distribution of inconsistently annotated cells, colored by cell type annotation from cell-morphology segmentation. **B.** Density of number of segmented cells across sample, using window of 400 x 400 µm. **C.** Density of number of segmented cells with discordant annotation between cell-morphology and nuclei-only segmentations, using window of 400 x 400 µm. **D.** Gene pair mutual exclusivity, based on premise that some genes do not typically co-exist in the same cell. Calculated as the proportion of cells expected to be mutually exclusive but observed to co-express both genes. X-axis is this proportion identified with nuclei-only segmentation, while y-axis is this proportion identified with a cell segmentation; either by cell-morphology (black), expansion-based (blue), or scRNA-seq (red). Only gene pairs with less than 1% co-expression in single cell data are shown. **E.** Cell type marker specificity of *MOBP* and *ARHGDIB*. Comparing relative presence of genes in different cell types across scRNA-seq, nuclei-only segmentation, cell-morphology segmentation, and expansion-based segmentation. For each cell type, the proportion of cells expressing the gene *MOBP* is reported as a positive score on the y-axis, while the proportion of cells expressing the gene *ARHGDIB* is reported as a negative score on the y-axis. Each segmentation strategy is depicted by a different shading. Excitatory and inhibitory neurons are combined into a single ‘neuron’ classification to align with the annotations of the *Ruiz-Moreno* dataset.

### Nuclei-only segmentations best preserve specificity of marker genes

Without a ground truth of cell type identity, it is difficult to assess the accuracy of the segmentation approaches with regards to correct cell type annotation. To remedy this issue bioinformatically, we investigated expected gene co-occurrence and mutual exclusivity. We first determined genes that seldom co-occur in cell types observed in glioma using a large single cell RNA sequencing (scRNAseq) reference dataset^16^. Compared to the scRNAseq dataset, mutually exclusive gene pairs had significantly greater co-occurrence spatially, the most frequent in expansion-based segmentation (**Figure 2d, Figure S2k**). For example, *MOBP* is a robust oligodendrocyte marker, with co-expression with *ARHGDIB,* an immune/vasculature marker, in less than 2% of cells in the scRNAseq dataset. By comparison, *MOBP* and *ARHGDIB* were co-expressed in 9% of nuclei-only segmentations, 23% of cell-morphology segmentations, and in 30% of expansion-based segmentations (**Figure 2e**). In addition to increased co-occurrence of mutually exclusive gene pairs, the specificity of cell type markers was considerably reduced. In ST data, *MOBP* was found in all cell types, and at higher proportions using expansion-based and cell-morphology segmentation masks, with similar findings for *ARHGDIB* (**Figure 2e**). Although annotations are largely concordant, expansion-based and cell-morphology segmentation introduced greater levels of transcript misassignment that have the potential to misinform annotation strategies. Hence, annotations based on nuclei-only segmentation are sufficient to provide annotations and best preserve properties of marker genes without significant loss in overall numbers of high-quality cells.

### Cell-morphology segmentation detects multinucleated cells

Due to their nature of cell identification based solely on nuclei, both expansion-based and nuclei-only segmentations are incapable of identifying multinucleated cells. Tumor tissue contains an increased abundance of multinucleated cells^17,18^ and cell-morphology segmentation is required for their exploration. In our glioma sample, 5% of segmented cells (2,028 cells) contained multiple nuclei (**Figure 3a**). However, cell-morphology estimations in areas of high cellular density can become inaccurate and contribute to false identification of multinucleated cells. To investigate whether cell-morphology segmentation identifies truly multinucleated cells, we conducted an enrichment analysis within each cell type. In line with expectations, this analysis confirmed that all highly proliferative tumor cell states except mesenchymal-like tumor cells^19^ were enriched for multinucleated cells (**Figure 3b**). This finding remained relatively stable when assessing only cells with three or more nuclei, as well as including cells containing multiple different annotations (**Figure 3b**). Of all identified multinucleated cells, 54% only contain nuclei with the same annotation, and 94% contained at least two nuclei with the same annotation. Considering all nuclei individually, 72% of nuclei in multinucleated cells matched the annotation derived from cell-morphology segmentation, compared to 89% for single-nucleated cells. Multinucleated cells containing multiple annotations represented 17% of all discordant annotations. The most common combination of multiple annotations was oligodendrocytes and excitatory neurons, indicating a potential failure to recognize the ensheathing nature of oligodendrocytes as distinct to the neuron itself (**Figure 3d**). Hence, discordant nuclei annotations in the same cell were associated with the healthy tissue region, while consistent annotations associated with the leading edge and a highly concentrated region in the tumor (**Figure 3c**). In summary, cell-morphology segmentation can identify truly multinucleated cells, allowing investigation into this key component of tumor architecture.

**Figure 3.**
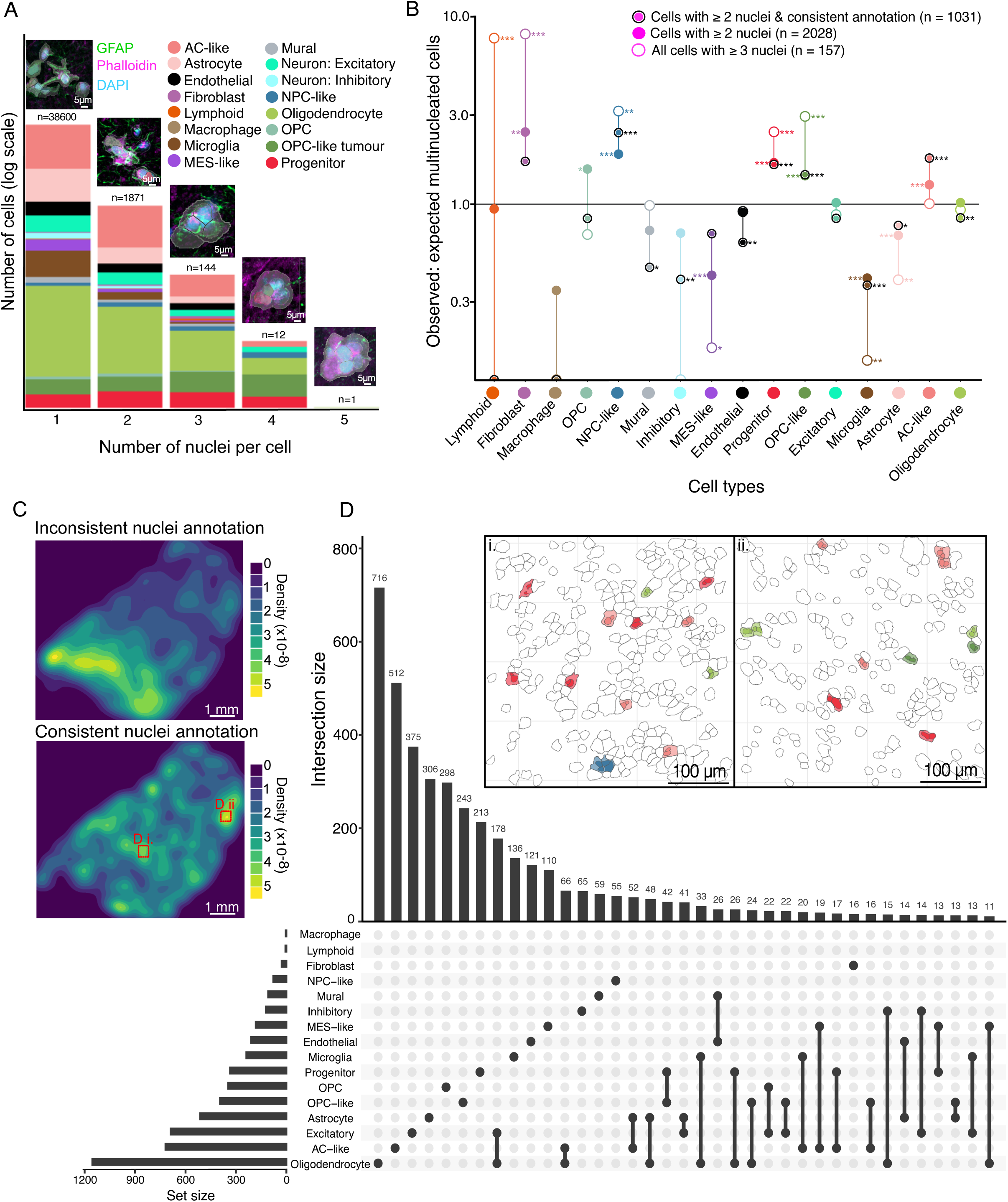
Cell-morphology segmentation enables the identification of multinucleated cells. **A.** Distribution of cells with different numbers of nuclei in sample, colored by relative proportion of each cell-type. Representative image of cell-morphology and nuclei-only segmentation overlaid with IF staining shown above for each number of nuclei. Scale, 5 µm. **B.** Over-abundance of multinucleated tumor cell types. Ratio of observed to expected number of multinucleated cells assuming equal likelihood of any cell type being multinucleated. Chi-squared test. Filled circles: all multinucleated cells, irrespective of annotation consistency (n=2,028); empty circles: all cells with 3 or more nuclei, irrespective of annotation consistency (n=157); bolded circles: all multinucleated cells with consistent annotation (n=1,031). Cell types ordered from left to right by overall abundance in sample; more abundant cell types require smaller ratio deviations from 1 to reach significance, and ratios of less abundant cell types are easily influenced by false positives. **C.** Density of multinucleated cells across sample. Top, multinucleated cells with inconsistent nuclei-only and cell-morphology annotations. Bottom, multinucleated cells containing nuclei with matching nuclei-only and cell-morphology annotations. Scale, 1 mm. Regions (i) and (ii) marked in red used as insets in (D). **D.** Upset plot showing frequency of nucleus annotation combinations in multinucleated cells. Insets i. and ii., representative regions highlighting distribution of multinucleated cells. Scale, 100 μm.

### Tumor neighborhoods are associated with distinct unified clusters driven by metabolism

By integrating ST and SM, we can elucidate how metabolic alterations may affect or be affected by the composition of cellular neighborhoods. Using STalign^12^, metabolite signals from a serial section (10 µm distance from ST slide) were aligned and combined with transcriptomic data via assignment to nearest cells as determined by nuclei-only segmentation **(Fig. 3a, Methods)**. Signal from both modalities was well aligned as cellular neighborhoods estimated from ST (**Figure 3b-c**) and unified clusters that integrate SM (**Figure 3d-e**) similarly demarcated the histologically annotated regions of the sample based on H&E assessment into the normal, leading edge, and tumor regions (**Figure S1a**). Through this analysis, we found that the enrichment of cell types in each unified cluster mirrored their proportions in spatially aligned cellular neighborhoods (**Figure 3f**). However, while considering both metabolites and transcripts simultaneously, the unified clusters were overwhelmingly dictated by metabolite expression rather than cell annotation, indicated by the heterogeneity of cell types present within each unified cluster (**Figure 3g-h**). This suggests that the untargeted panel of metabolites drives differences beyond cell annotation, which the 10x Genomics Xenium panel is powered to detect, and can identify more nuanced neighborhoods in the spatial data.

### Unified multi-omic analysis reveals invasive biology at the tumor leading edge

Next, to detect whether integrated spatial multi-omic data allowed investigation of glioma tumor biology across molecular levels, we investigated the characteristics at the tumor leading edge. This region was characterized by the presence of actin-positive protrusions stretching across the tumor and into the normal tissue, detected by IF staining (**Figure 5a-c**). Altered expression patterns were investigated in cell neighborhoods unique to the tumor leading edge (N6 and N7) to explore this under-characterized architecture (**Figure S3**). Compared to the tumor core, multinucleated and OPC-like cells were 1.1 and 1.65-fold enriched in the leading edge, respectively (**Figure 3c**, **4f**). Differential expression analysis of OPC-like cells identified the upregulation of *PDGFRA* and *EGFR* in the leading edge (N6 and N7) compared to OPC-like cells in the normal tissue (N3) and tumor core regions (N8 and N9) (**Figure 5d, Figure S4a-d**). Both *PDGFRA* and *EGFR* play crucial roles in cell proliferation and migration^19–21^. Although these genes were also enriched in the sparsely localized AC-like tumor cells of the leading edge, they were significantly more highly expressed within the abundant OPC-like population. Interrogation of the SM data in the leading edge (U4) compared to both normal tissue (U1) and tumor core regions (U5 and U6) identified an abundance of glycerophosphocholine, a breakdown product of cell membrane phospholipids^22–24^ occurring during cellular turnover and migration (**Figure 5e, Figure S3e-f**). This suggests that OPC-like cells in the leading edge may play an active role in the infiltration of glioma into the normal brain. Overall, observations derived from ST and SM at the leading edge suggest highly congruent biological findings.

**Figure 4.**
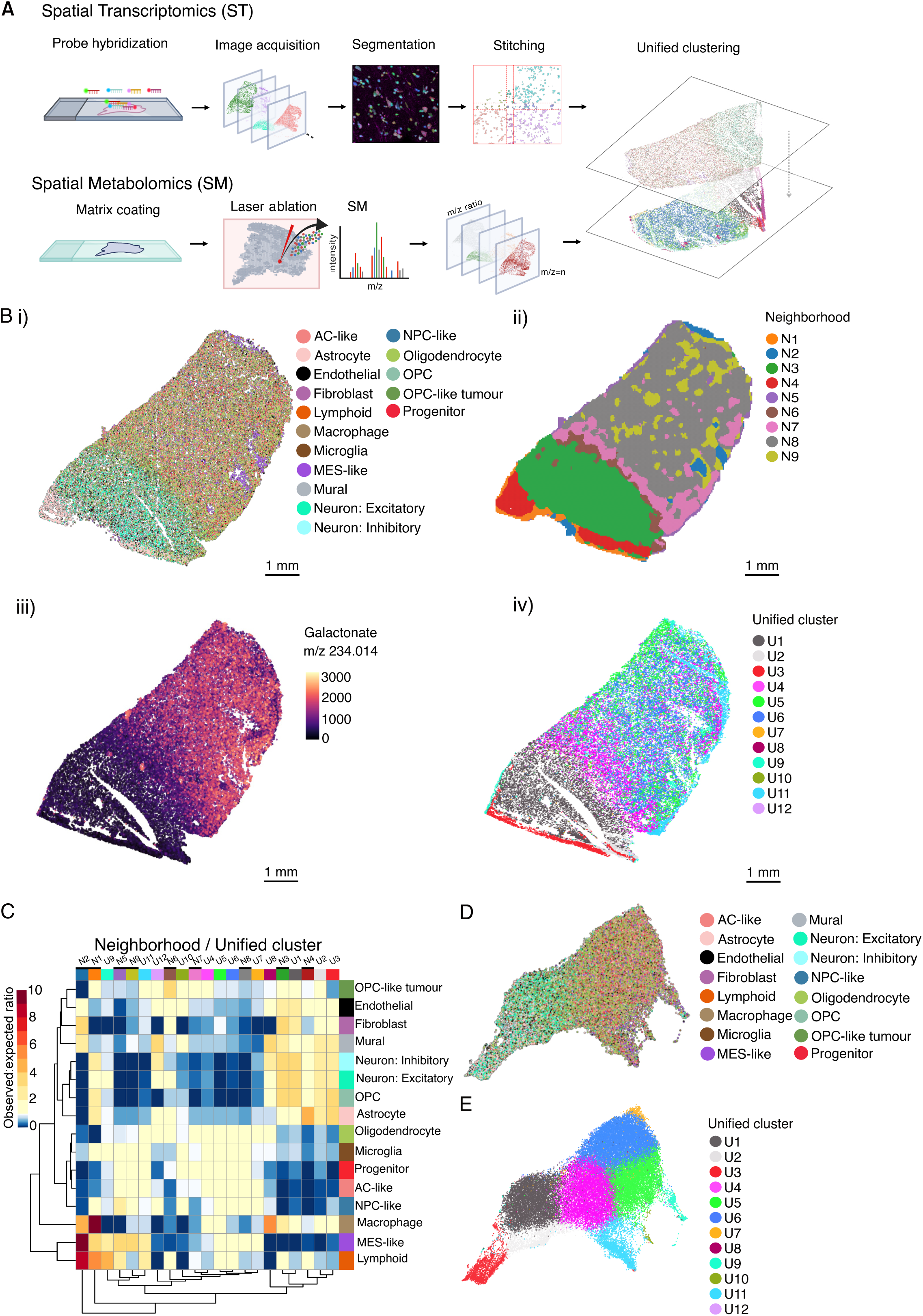
Tumor neighborhoods are associated with distinct unified clusters. **A.** Overview of preprocessing and alignment of spatial modalities. **B.** Spatial distribution in the common coordinate framework of i) annotated cells based on nuclei-only segmentation, ii) cellular neighborhood distribution based on cell type abundances in 160 x 160 µm windows, iii) spatial distribution of galactonate (m/z 234.0145) abundance, localized to each cell and iv) spatial map of unified clusters. Scale, 1mm. **C.** Relative abundance of cell types in each cellular neighborhood (N) and unified cluster (U). Only clusters with over 70 cells were included. **D.** UMAP derived from unified metabolomic and transcriptomic data, colored by nuclei-only cell type annotation. **E.** UMAP colored by unified clusters.

**Figure 5.**
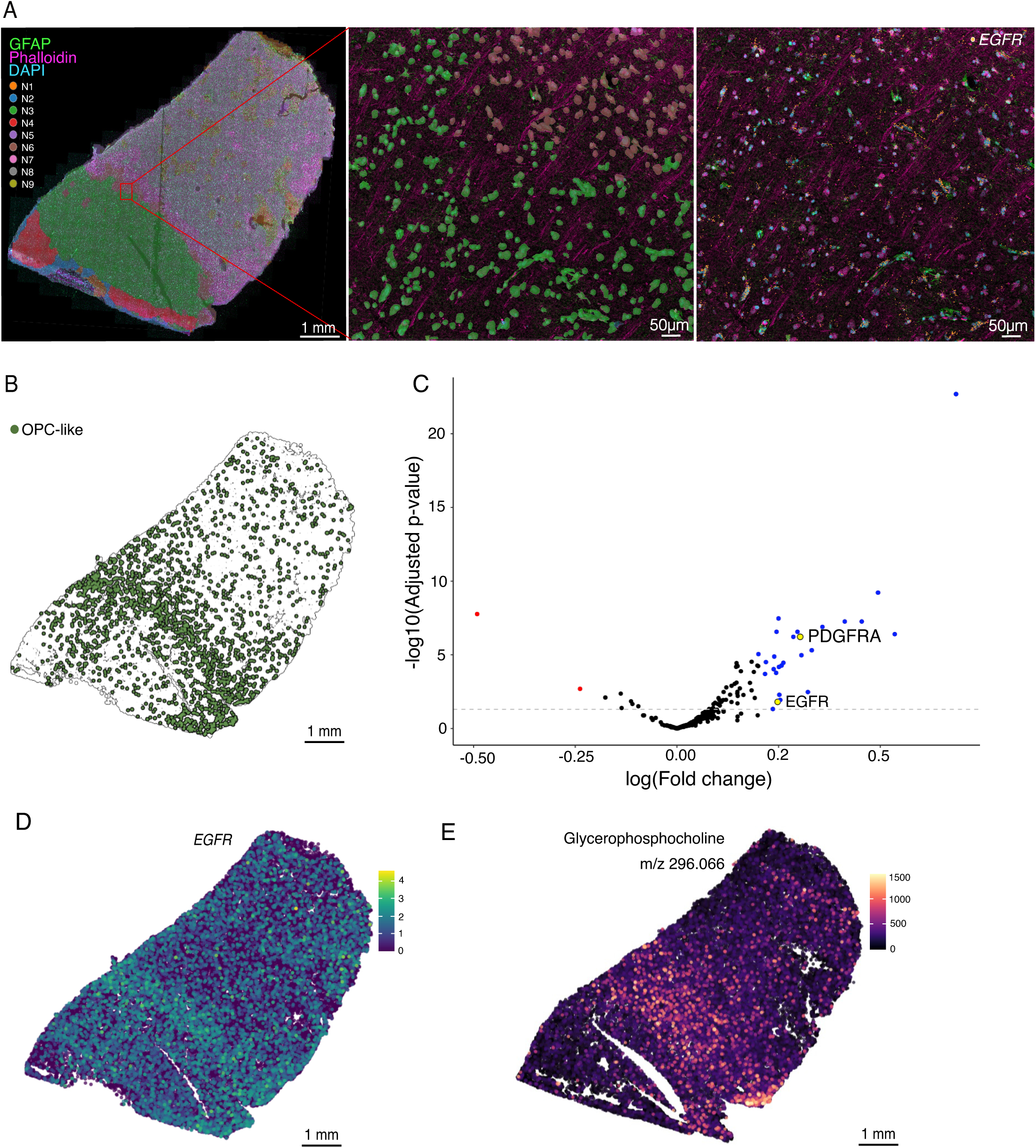
Unified analysis of the glioma leading edge. **A.** Spatial expression of *EGFR* transcript at the leading edge. Left, overlay of cell neighborhoods and IF staining. Glial fibrillary protein (GFAP, green), actin (Phalloidin, magenta) and nuclei (DAPI, cyan). Scale, 1mm. Middle inset, overlay of cell masks corresponding to cell neighborhoods N6 (brown) and N3 (green). Scale, 50 µm. Right inset, *EGFR* transcript localization (orange points) and IF staining. Scale, 50 µm. **B.** Distribution of OPC-like tumor cells across sample, annotated by nuclei-only segmentation. **C.** Volcano plot of differentially expressed genes in OPC-like tumor cells at the leading edge, compared to OPC-like tumor cells located in normal or tumor core tissue regions. Dotted line indicates significance. **D.** Spatial distribution of *EGFR* and *PDGFRA* gene expression. Scale, 1 mm. **E.** Spatial abundance of Glycerophosphocholine (m/z:[M+K]-296.066). Scale, 1 mm.

## Discussion

Here we present Spatial Multi-omics Integration (SMINT), a workflow to combine ST and SM data that addresses the main challenges faced in the integrative analysis of spatial multi-omics data: cell segmentation and alignment (**Fig. 6**). The SMINT workflow consists of 1) conducting ST and SM on serial sections 2) immunostaining ST slides for bespoke segmentation 3) cell type annotation using nuclei-assigned transcripts 4) registration of ST and SM datasets to a common coordinate framework and 5) integration of ST and SM data for unified clustering at cellular resolution. By combining existing methods with custom algorithms, the SMINT workflow yields a comprehensive, high-resolution representation of cellular and molecular organization within the heterogeneous glioma microenvironment.

**Figure 6.**
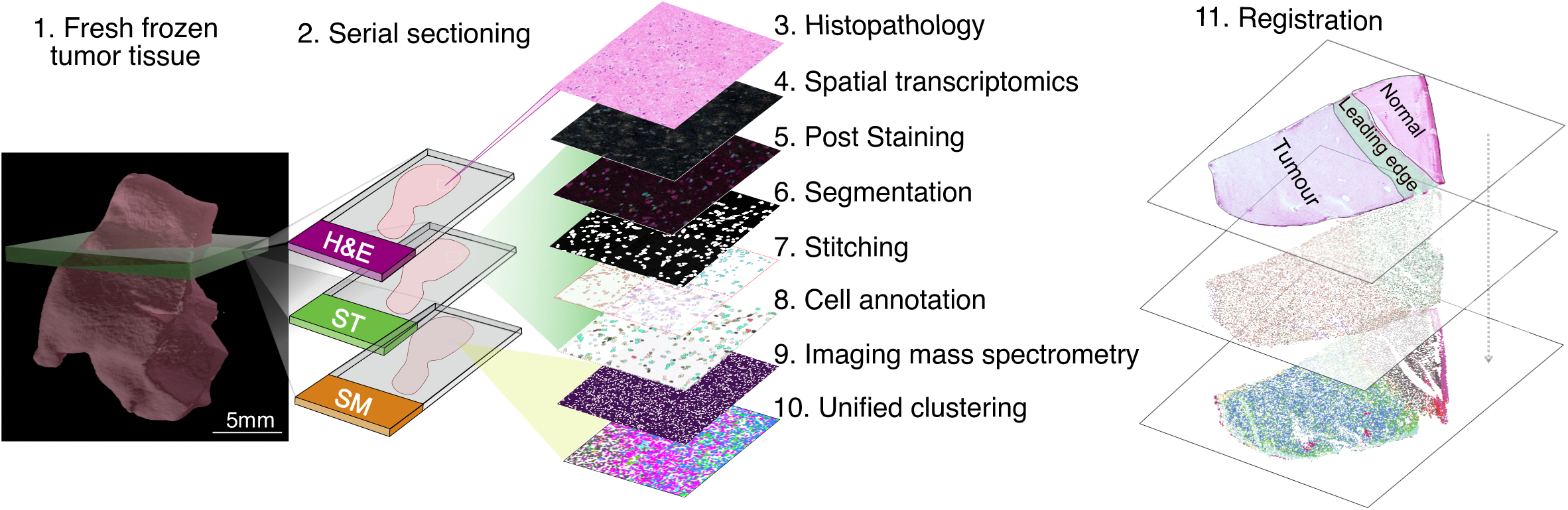
Summary of Spatial Multi-omics Integration (SMINT) workflow.

Our comparison of different segmentation strategies suggests that nuclei-only segmentation suffices, when multinucleated cells are not of interest. This finding is significant as it suggests that nuclei-only segmentation can serve as a reliable alternative to more computationally intensive methods, particularly in contexts where rapid and scalable analysis is required. Moreover, the ability to achieve high concordance with scRNAseq suggests that nuclei-only segmentation reduces the risk of transcript misassignment, which is a common issue in two-dimensional segmentation methods. In particular, the sensitivity and specificity of cell type specific genes was found to be especially compromised in cell-based approaches when compared to scRNAseq data. This possibility was first highlighted by Littman *et al.* (2021)^25^ when assessing misassignment of transcripts due to segmentation errors in MERFISH results of mouse hippocampal samples where spatial gene expression patterns were already well defined. Our findings further support these observations in samples that have not been spatially characterized before. Furthermore, differential expression analyses are significantly hampered by molecular contamination in ST, presenting nuclei-only segmentations as a favorable alternative to cell-based methods^26^.

It is important to note that nuclei-only segmentation does not limit all downstream analysis to nuclei-bound transcripts. Cell-free methods of analyzing expression are becoming increasingly utilized across all steps of transcriptomic processing, including differential gene expression analysis^27^ and neighborhood detection^28,29^. These methods pair with cell type annotations based on nuclei-only segmentation and can enhance tumor characterization, offering a flexible foundation from which pipelines tailored for specific biological hypotheses can be developed. Nonetheless, the inability to correctly quantify cell type numbers with these methods advocates for a segmentation strategy in tumors.

Using this integrated workflow in glioma, we revealed regional distinctions in the tumor composition. In the tumor, bulk cell type composition did not dictate metabolite activity, highlighting the intricate relationship between spatial organization and metabolic abundance. This complexity is particularly pronounced at the tumor leading edge. We found that the leading edge was enriched for OPC-like cells, which exhibited upregulated growth factor receptor expression (*EGFR* and *PDGFRA*), coexisting with increased glycerophosphocholine abundance. We therefore speculate that the glycerophosphocholine, produced by enhanced membrane turnover and remodelling, is driven by OPC-like cell-mediated invasion. This is further supported by the presence of actin-positive protrusions, indicative of white matter tracts, stretching across the leading edge from tumor into normal tissue. Clinical observations suggest that glioma preferentially invade along these white matter tracts^30–32^, despite evidence that white matter serves as an inhibitory substrate for neurite outgrowth and astrocyte migration^33,34^. When taking into account the highly motile nature of OPC-like cells^35,36^, our results suggest they may utilize white matter tracts as a scaffold to mediate invasion, facilitating their progression and contributing to the invasive characteristics observed in gliomas.

In conclusion, the approach presented here allows for a nuanced understanding of tumor heterogeneity and the microenvironment and has the potential to uncover novel biological insights that would be obscured if analyses were conducted separately.

## Limitations

A main limitation of our workflow is the computational rigor and associated resources required for cell segmentation and alignment. Cell segmentation at maximal resolution requires uncompressed file scans necessitating the introduction of stitching cell masks, which is especially computationally burdensome. Similarly, alignment with single cell precision is computationally demanding but necessary to ensure modalities can be integrated. Notably, it is possible to conduct both ST and MALDI-IMS on the same section without requiring multiple rounds of alignment^37^. However, conducting analyses on separate slides enhances the quality of the data by allowing optimal quality for the generation of each modality.

While our findings with regards to segmentation strategies and multinucleated cells are based on a single sample, the features of glioma are likely to make these findings more generalizable. Similarly, our findings of altered choline metabolism validated across multiple modalities is demonstrated for a single sample only and requires further exploration.

## Resource Availability

### Lead contact

Further information and requests for recourses and reagents should be directed to and will be fulfilled by the lead contacts, Saskia Freytag and Sarah Best.

### Materials availability

This study did not generate new unique reagents.

### Data and code availability

All code for replicating analysis can be found in our GitHub page (https://github.com/JurgenKriel/SMINT).

## Acknowledgements

We thank A. Fakhri, S. Stylli and K. Drummond for expert curation of the Royal Melbourne Hospital Neurosurgery Brain and Spine Tumour Tissue Bank, the WEHI Histology core, C. Anttila for spatial transcriptomics assistance at the WEHI Cellular Genomics facility and P. Rajasekhar for image analysis consulting.

This work was made possible and financially supported in part through the authors’ membership of the Brain Cancer Centre, support from Carrie’s Beanies 4 Brain Cancer, the Venture Grants Scheme administered by Cancer Council Victoria (VG2022 to S.A.B., S.F., J.R.W. and M.J.M.) and through Victorian State Government Operational Infrastructure Support and Australian Government NHMRC Independent Research Institutes Infrastructure Support Scheme (IRIISS). Support from the Victorian Cancer Agency Mid-Career Research Fellowship (MCRF22003 to S.A.B.), WEHI Johnson PhD Scholarship and an Australian Government Research Training Program (RTP) Scholarship (J.J.D.M.), CSL Translational Data Science Scholarship (J.J.D.M.), WEHI IPSI Scholarship (O.E.F.), Melbourne Research Scholarship (O.E.F.) and Gregg Symons Scholarship for Brain Cancer Research (O.E.F.).

## Author contributions

J.J.D.M., J.K., S.F., S.A.B. and J.R.W. conceived the study. A.V. and A.M. processed the patient tissue, V.K.N. and M.J.M. performed MALDI-TOF imaging, J.K., J.J.D.M. and T.Y. processed the raw data, J.J.D.M. and T.Y. performed the bioinformatic analysis, J.K. performed imaging analysis and O.E.F. performed the pathology analysis. S.F., S.A.B. and J.R.W. obtained funding and provided supervision. J.J.D.M., J.K., S.F., S.A.B. and J.R.W. wrote and revised the manuscript with input from all authors.

## Declaration of interests

The authors declare no competing interests.

## STAR Methods

### Key Resource Table

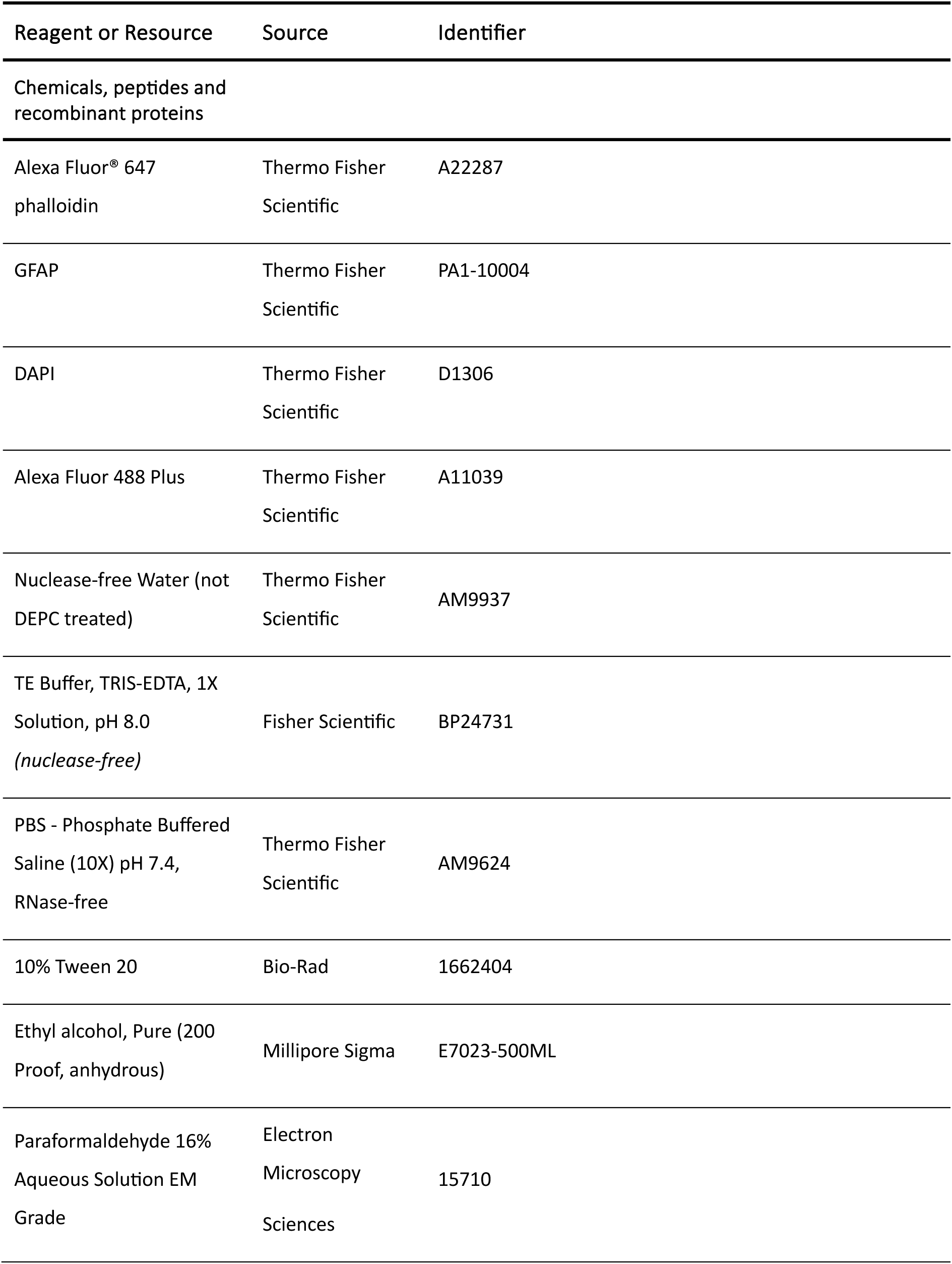
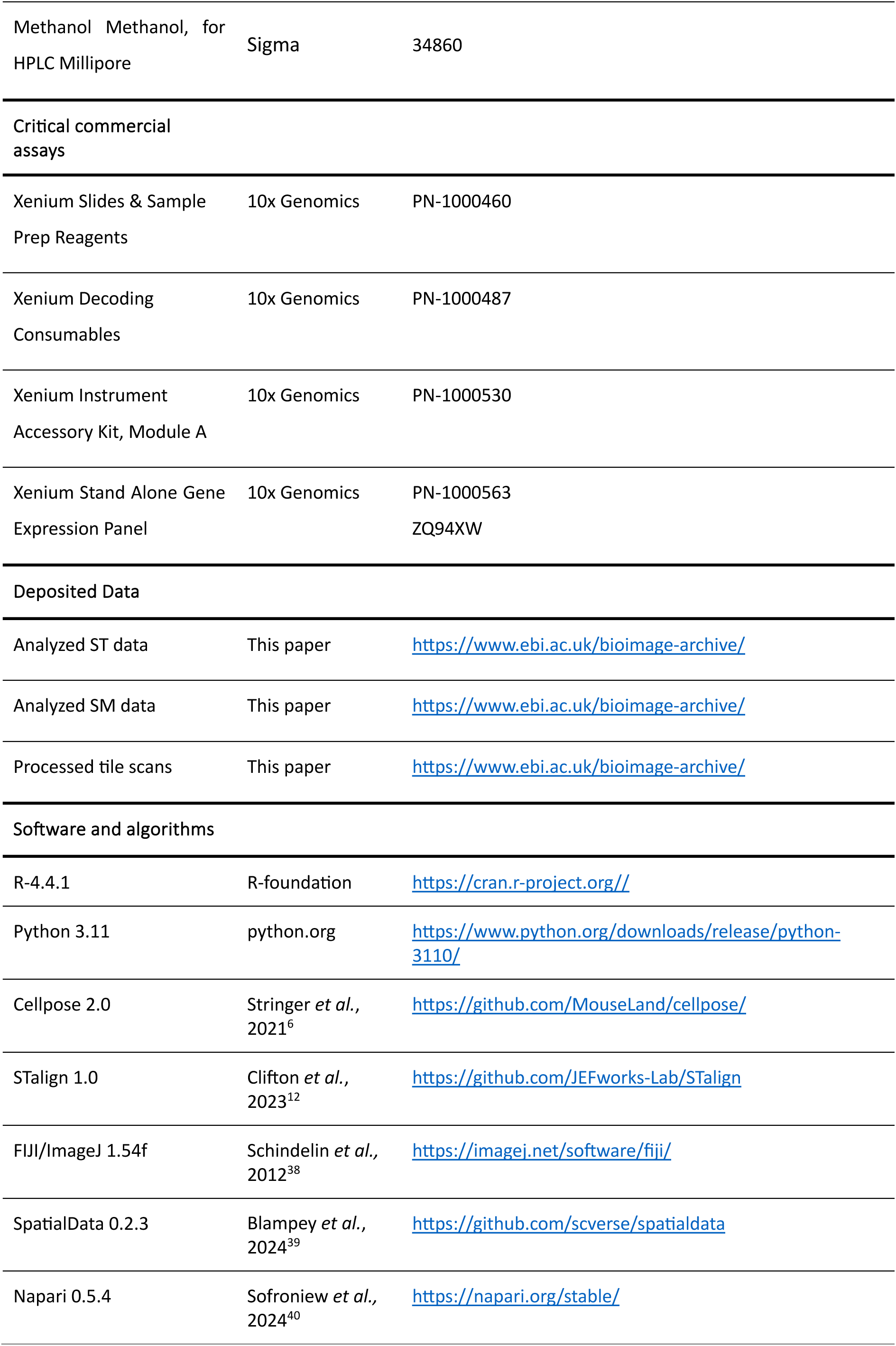
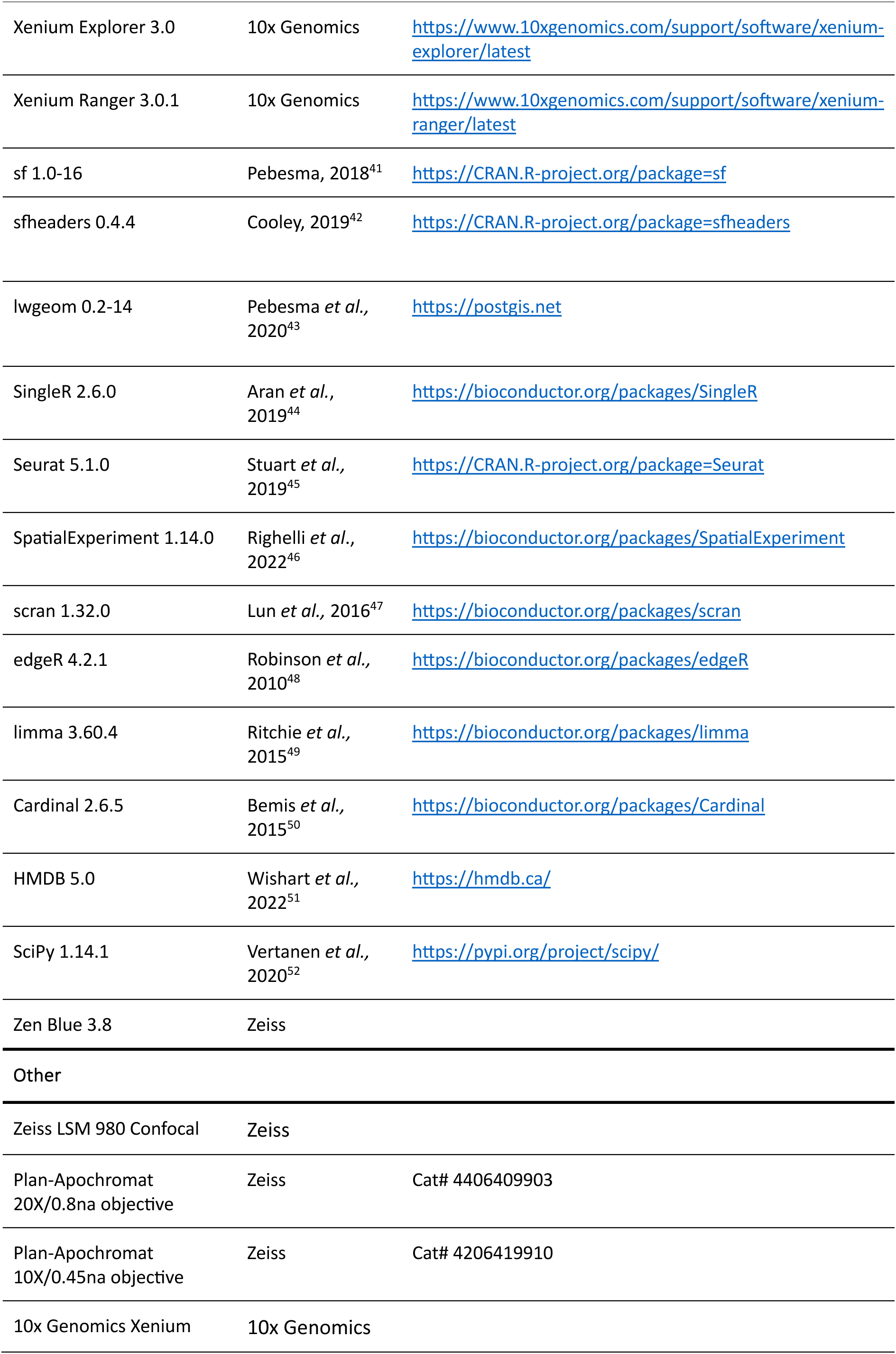
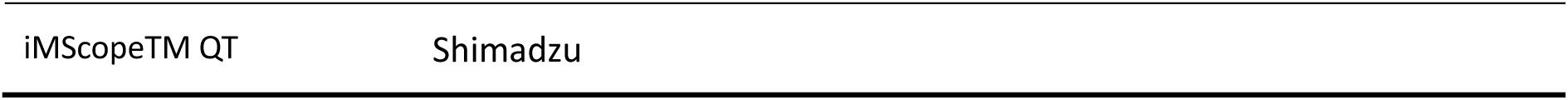

### Experimental Model and Study Participant Details

#### Human subjects

One de-identified astrocytoma WHO CNS grade 3 glioma sample was obtained from the Royal Melbourne Hospital Neurosurgery Brain and Spine Tumour Tissue Bank (Melbourne Health Ethics # 2020.214) and analyzed under a protocol approved by the Walter and Eliza Hall Institute Human Research Ethics Committee (HREC 21/21).

### Methods Details

#### Fresh-frozen histology

Patient samples were obtained directly from surgery via the Royal Melbourne Hospital Neurosurgery Unit and were immediately flash-frozen directly in isopentane in a liquid nitrogen bath. Frozen tissue was stored at − 80 °C until downstream sample preparation. Frozen tissues were sectioned using a Cryostar NX70 at −20 °C. Briefly, samples were mounted, equilibrated to temperature and trimmed to the region of interest. Once the region of interest was confirmed, serial sections without OCT were cut at 10 µm and collected onto each respective series of slides. Slides and samples were kept cold throughout sectioning, to minimise freezing and thawing cycles.

To assess the quality of sections, representative slides were thawed, formalin fixed and stained with Haematoxylin and Eosin (Leica AutoStainer, ST5010).

#### 10x Genomics Xenium in situ gene expression

10 μm thin frozen sections were cut onto 10x Genomics Xenium slides. Tissue sections were first fixed using 4 % paraformaldehyde for 30 minutes at room temperature. Following fixation, the slides underwent permeabilization consisting of a 1-minute wash in 1X PBS, followed by a 2-minute incubation in 1 % SDS, and additional 1-minute washes in 1X PBS. The slides were then incubated in pre-chilled 70 % methanol for 60 minutes on ice to further permeabilize the tissue, then washed twice more in 1X PBS for 1 minute each.

Custom probe panels were hybridized to the target RNA overnight at 50 °C using 34 μl of probes per slide, with the Xenium Probe Hybridization Buffer (PN-2000390) and TE Buffer. Following hybridization, unbound probes were removed through washes with PBS-T, prepared by mixing 1X PBS (prepared from 10X PBS, pH 7.4) and 0.05% Tween-20, to reduce non-specific binding.

After washing, the ligation process was performed using the Ligation Mix, which included 481.2 μl of Xenium Ligation Buffer (PN-2000391), 13.8 μl of Xenium Ligation Enzyme A (PN-2000397), and 55.0 μl of Xenium Ligation Enzyme B (PN-2000398) per reaction. Circularized probes were enzymatically amplified using the Amplification Master Mix, which consisted of 495.0 μl of Xenium Amplification Mix (PN-2000392) and 55.0 μl of Xenium Amplification Enzyme (PN-2000399) per reaction. Autofluorescence quenching was conducted using 500 μl of Autofluorescence Solution, prepared with Xenium Autofluorescence Mix (PN-2000753) and 100 % ethanol, to reduce background noise. Nuclei were stained with 500 μl of Xenium Nuclei Staining Buffer (PN-2000762) to aid in tissue identification during imaging. The slides were then loaded into the Xenium Analyzer for imaging and analysis. Throughout the protocol, reagent handling and storage conditions were maintained, including the use of TE Buffer for probe resuspension and specific thermal cycler settings, to ensure assay performance.

#### Post-staining

For post-staining 10x Genomics Xenium slides, no additional fixation or permeabilization was required. Slides were incubated with blocking buffer (0.1 % Tween-20 in 1x PB with 5 % goat serum) for 1 hour at 4 °C. After washing the slides three times with PBS-T, primary antibodies, diluted in the blocking buffer, was applied and incubated overnight at 4 °C. The slides were then washed three times with cold PBS-T before adding the secondary antibodies, which was diluted at 1/1000 in the blocking buffer and incubated for 1 hour at room temperature. Following another set of three washes with PBS-T, DAPI was added at a 1:1000 dilution and incubated for 10 minutes at room temperature. The slides were washed three more times with PBS-T, mounted with FluoroMount solution, and stored at 4 °C until imaging.

#### Image acquisition

##### Stage calibration

In order for the LSM 980 confocal stage to recognize the 10x Genomics Xenium slide as a conventional 76 x 26mm slide, support points were assigned to each corner of the 10x Genomics Xenium slide sample area for stage calibration. The Zen Blue experiment file was reused for subsequent rounds of imaging.

##### Set regions of interest and support points

A low resolution TPMT tilescan (256×256 pixels) of the entire sample area was taken at 10x magnification. Regions of interest were drawn accordingly for each section on the sample area. To ensure all tiles were captured at adequate focus, 10-20 support points were added to each section depending on their size.

##### Setting imaging parameters

Using a 20x objective, the appropriate strength of 405, 488 and 647 laser lines was determined. Thereafter, the z-position of each support point was calibrated manually, opting for a z-slice in the middle of the section. Tile sizes were set to 3852 x 3852 with a LSM scan speed of 7 at bidirectional direction. Small z-stacks of 5-6 slices were selected to reduce imaging time. After determining the upper and lower z-limit, the center slice was determined, and the z-stack strategy set to center for tile scan acquisition.

#### Image processing and export

Orthogonal projections were first created using Zen Blue software (version 3.8). Thereafter, individual tiles were exported as tiffs, followed by stitching of the orthogonal projection in Zen Blue at 10% overlap. Resulting cell masks after segmentation were overlaid with the stitched output.

#### Training a custom Cellpose model

Pre-processing of images required merging of the 488 and 647 channels (FIJI/ImageJ, version 1.54f), to ensure that cell outlines of GFAP negative cells would still be identified. 50 tiles from various regions of the original tile scan were then imported into Cellpose (version 2.0) and an initial segmentation run was conducted using the pretrained Cellpose model. The settings were as following *Flow Threshold*: 1, *CellProbability Threshold*: 0.4, *Cell Diameter:* 100, *Channel to segment*: 1 (the merged 488 and 647 images), *Additional Channel to segment:* 2 (405 Dapi channel).

These initial masks were then assessed on a per tile basis, with inaccurate segmentations deleted and manually redrawn for each cell. The segmentation files were then exported as seg.npy files for training. Training of the Cellpose model was run on a Nvidia A30 GPU at a learning_rate of 0.1, with a weight_decay of 0.0001 and 1000 epochs.

#### Cell segmentation

Segmentations were run on all exported tiles using the Cellpose CLI segmentation function. For final segmentation, the cellpose flow_threshold was set to 2.0, cellprob_threshold to −1 and diameter to 100.5. Outline coordinates were then saved as a text file for subsequent stitching in R, along with seg.npy files for viewing. The pretrained Cellpose nuclei model was used for nuclei-only segmentation.

#### Stitching

Stitching was performed in R separately and identically for nuclei and cells, using plugins for Simple Features (packages: sf version 1.0-16, sfheaders version 0.4.4, and lwgeom version 0.2-14). First, a single object combining segmentations from all tiles was created, with tiles localized to their global position. Conflicting polygons were identified in tiled regions and were assessed for the proportion of area of each polygon within their overlapped area. If the maximum overlap was greater than 50%, polygons were combined; if the maximum overlap was less than 25%, the overlap was removed from each polygon, and if the maximum overlap was between these values, the smaller polygon was removed. Of all polygon conflicts, only 191 polygons (0.03%) were completely removed for nuclei, and 190 (0.03%) for cells. 99.6% of total nuclei area and cell area was retained (for a complete summary of procedures and changes to nuclei and cell number see **Figure S1**).

Cells and corresponding nuclei were matched using a similar framework. Nuclei were assessed for their overlap with segmented cells – both area and number of cells. 80% of overlapping nuclei overlapped with a single cell. Of these, 98% were completely encapsulated. For the rest, if 50% or more of a nucleus area was contained within a cell and the rest is outside of other cells, the additional nucleus area was added to the cell-morphology segmentation. If one had less than 50% of its shape within a cell, the cell was removed. In the case of nuclei overlapping multiple cells, the nucleus was divided based on the cell boundaries and divisions were kept if enough of its area was contained by the individual cell - 40% for overlaps with 2 cells and 30% for overlaps of 3 or more - otherwise the nucleus region in question was removed. If cells finished with no matched nuclei, they were removed. If nuclei had no matched cell, the nucleus segmentation was also used as the cell-morphology segmentation.

#### Transcript alignment

To register transcript locations to nuclei-only and cell-morphology segmentations, affine transformation was applied on the centroids of the post-stained nuclei, aligning them to the centroids of the 10x Genomics Xenium nuclei. Cell-morphology segmentation was then registered by applying the same transformation as the nucleus centroid closest to their own. Assigning transcripts to segmentations was performed using the st_join function of the sf package (version 1.0-16). Registration to transcripts requires micrometer precision, and the increased density of transcripts with segmented nuclei vs cytoplasm indicates this was achieved (0.012 transcripts per µm in nuclei vs 0.06 transcripts per µm in cytoplasm).

Any nuclei with less than 20 transcripts or less than 5 UMIs were subsequently removed, along with any cells with no nuclei passing this threshold (19% of cells with ≥ 20 transcripts). Additionally, cells with more than 3 negative control probes or 3 unassigned probes were removed along with their bound nuclei. Reads from negative control or unassigned probes were subsequently removed before downstream analysis. These same thresholds were also applied to the expansion-based segmentation.

#### Annotation

Annotations were performed separately across nuclei-only and cell-morphology segmentations according to *Moffet et al.*^53^ enlisting semi-supervised annotation followed by manual refinement. Counts were log-normalized and clustered via PCA based on the first 25 principal components. Individual cell annotations were estimated using SingleR (version 2.6.0) and two references; *Ruiz-Moreno et al.*^16^ to identify non-neoplastic populations, and *Couturier et al.*^15^ to characterize the neoplastic characteristics. A UMAP projection was used to visualize the cell populations based on the first 25 principal components with default parameters, excluding min_dist, which was set to 0.1. Each cluster was preliminarily assigned the most common annotation within the cluster. Clusters were then confirmed or modified based on overexpression of marker genes. These annotations were performed cyclically, first to define larger populations (tumor, immune, vasculature, glial, neuronal), then reperforming clustering on these populations separately to pull out more specific annotations.

#### Niche analysis

Cell neighborhoods were determined as in *Moffet et al*.^53^ Briefly, 160 µm wide windows were laid across the sample with 40 µm shift, and cell types within each window were tallied. Windows were clustered into 9 neighborhoods based on cell type abundances using a distance metric, and each 40 µm square was assigned the most abundant cluster across the 16 windows overlapping the region. Cells were assigned the neighborhood overlapping their centroid location.

#### Spatial metabolomics (SM)

Frozen tissue was sectioned at a thickness of 10 µm directly onto Indium Tin Oxide (ITO) coated glass slides (surface resistance of 70-100 Ω). Frozen sections were dried in freeze dryer (MODULYOD, Thermo Electron Corporation) for 30 min, followed by collection of optical images using the light microscope embedded in the MALDI-TOF MSI instrument (iMScope^TM^ QT) prior to matrix application.

Matrix deposition with α-cyano-4-hydroxycinnamic acid (CHCA, P# C2020, Sigma-Aldrich) by sublimation (0.7 µm thickness at 250 °C) using iMLayer (Shimadzu, Japan), followed by CHCA matrix spraying using iMLayer AERO (Shimadzu, Japan) to recrystallize and obtain fine matrix crystals, that enable high sensitivity and high spatial resolution. For CHCA matrix spraying, 8 layers of 10 mg/mL CHCA in acetonitrile/water (50:50, v/v) with 0.1 % trifluoroacetic acid solution were used. The stage was kept at 70 mm/sec with 1 sec dry time at a 5 cm nozzle distance and pumping pressure kept constant at 0.1 and 0.2 MPa, respectively.

All SM experiments were performed using iMScope^TM^ QT instrument (Shimadzu, Japan). The instrument is equipped with a Laser-diode-excited Nd:YAG laser, and an atmospheric pressure MALDI. Data was acquired using a laser intensity at 60 for 10 µm spatial resolution, detector voltage was set at 2.36 kV, laser repetition frequency set at 1000 Hz, desolvation line temperature maintained at 250°C and a laser irradiation count of 50 shots were accumulated per pixel.

Data analysis was conducted as described in *Lu et al*.^13^ Briefly, data were imported into R and analyzed with Cardinal (version 2.6.5), including normalization via total ion current (TIC), spectrum smoothing, baseline correction, peak picking, alignment and filtering. Note that peaks that were also found in matrix-only sample preparations (no tissue) on a minimum detection frequency in at least 5 % of the pixels were also removed.

Next, the processed spectrum was subjected to binning to obtain the final spectrum information. Peaks were matched against the Human Metabolome Database (HMDB, version 5.0) by using the k-nearest neighbor methodology. For each peak, a set of 30 nearest neighbors from the generated HMDB peak list, for which the mass is less than 50 ppm distant, is identified.

For effective visualization of the intensity of any peak, we employed the application of a Gaussian Kernel Density Estimation (KDE) was used. The computation of the KDE was performed in R by using the stats::density (version 4.3.0) function.

The primary SM data was converted into coordinate-labeled data in a grid-like matrix format. To determine the boundary coordinates for both spatial images, the scipy.spatial.ConvexHull function (SciPy version 1.14.1) was applied. A rigid transformation was then performed on the SM coordinates to maximise the overlap with the ST data. Following this, the linear and translational components of the transformation were applied to the non-convex hull SM points, generating the final SM dataset, which is now ready for detailed registration.

#### Alignment of spatial metabolomics (SM) to spatial transcriptomics (ST) data

Cell centroid coordinates determined from stitched polygons were imported into python as numpy arrays. Metabolite coordinate information was transformed to numpy arrays for x and y coordinates separately. The STalign python package (version 1.0) was used for alignment, which relies on an iterative gradient descent to align spatial datasets. Target SM and source ST datasets were aligned at a similar angle using the pre-alignment.py script provided in our github, with theta_deg determining the angle of rotation. Cell centroid positions were then rasterized into density images using the STalign.rasterize function. To make these compatible with the np.imshow function, the STalign.extent_from_x function was applied to return a 4-tuple value to determine the extent of each image.

Landmarks for registration were determined manually using the curve_annotator.py module provided by STalign developer. Rasterized images were saved as .npz files to load into the point annotator. Coordinates were loaded as numpy arrays to be used as landmarks for diffeomorphic metric mapping through the STalign.LDDMM function. This jointly estimates an affine transformation as well as a diffeomorphism value between the two images. Once determined, these values were applied to the SM dataset using the STalign.transform_points_source_to_target function. The transformed x,y coordinates could then be exported for cluster analysis.

SM results were not originally assigned to cells and had a resolution of 10 µm. After registration of ST and SM slides, metabolite signals were assigned to cells by averaging the expression of each metabolite across all positions within 20 µm of the cell centroid, to ensure best coverage of cell-localized metabolites. These metabolite expression signals were combined with the log-normalized gene expression and underwent PCA, clustering and UMAP as previously described for cell type annotation.

### Quantification and Statistical Analysis

#### Morphometric analysis

Polygon areas were calculated using the sf package. Matching expansion-based segmentation to cell-morphology segmentation was possible due to alignment of post-stained polygons to Xenium transcript data, which is aligned to the expansion-based segmentation. Using the sf::st_intersection and Sf::st_area functions, the identity of overlapping polygons and the extent of their overlap relative to their original area were measured and retained if above 45% overlap. This resulted in 95% of cells to be considered truly matched.

Cell roundness was assessed by comparing the ratio of the radii of the maximum inscribed circle (lwgeom::st_cast, lwgeom::st_dist, lwgeom::st_buffer, lwgeom::st_circle_center) and minimum circumscribing circle (sf::st_minimum_bounding_circle) – values closer to 1 indicate a more circular shape. Number of neighbors within 20 µm of the cell centroid were recorded. Due to differences in segmentation strategies, this number may be different for expanded cells and their matched cell-morphology counterpart. To accommodate this, the average number of neighbors between the two was recorded.

#### Assessment of annotation concordance

The single cell dataset from *Ruiz-Moreno et al.*^16^ was downsampled cells to 227,136 cells across all annotations, 20 % of the total 1,135,677 cells. For all pairs of genes in the spatial sample and *Ruiz-Moreno* dataset, the number of cells with both genes present and compared to the number of cells with either gene present was calculated to assess how consistently these genes co-occur. This was also performed for the three morphology-based segmentations discussed in this paper.

#### Overrepresentation analyses

Overabundance of cell types in annotations using different segmentation strategies was calculated by Chi-squared tests with 2 degrees of freedom, performed individually for each cell type with global proportions of segmented nuclei to cells considered.

Overabundance of tumor cell states being multinucleated was calculated by Chi-squared tests with 2 degrees of freedom, performed individually for each cell type with global proportions of multinucleated cells considered.

Overabundance of cell types in niches or metabolites clusters were calculated separately by Chi-squared tests with 2 degrees of freedom, performed individually for each grouping with global proportions of cluster or niche affiliation considered.

#### Differential expression analysis

Signatures upregulated at the border were determined via differential gene expression Using edgeR (version 4.2.1) and subsequent KEGG pathway enrichment analysis using limma (version 3.60.4), with OPC-like cells within neighborhoods N6 and N7 assigned to ‘leading edge’, and the combined population of neighborhoods N3, N8 and N9 assigned to ‘comparison’. For metabolite signatures at the border, unified cluster U4 was assigned to ‘leading edge’ and clusters U1, U5 and U6 as ‘comparison’.

## Notes

### Competing Interest Statement

The authors have declared no competing interest.

