## Supplementary Figures for "An integrative spatial multi-omic workflow for unified analysis of tumor tissue"

Supplementary Figure 1

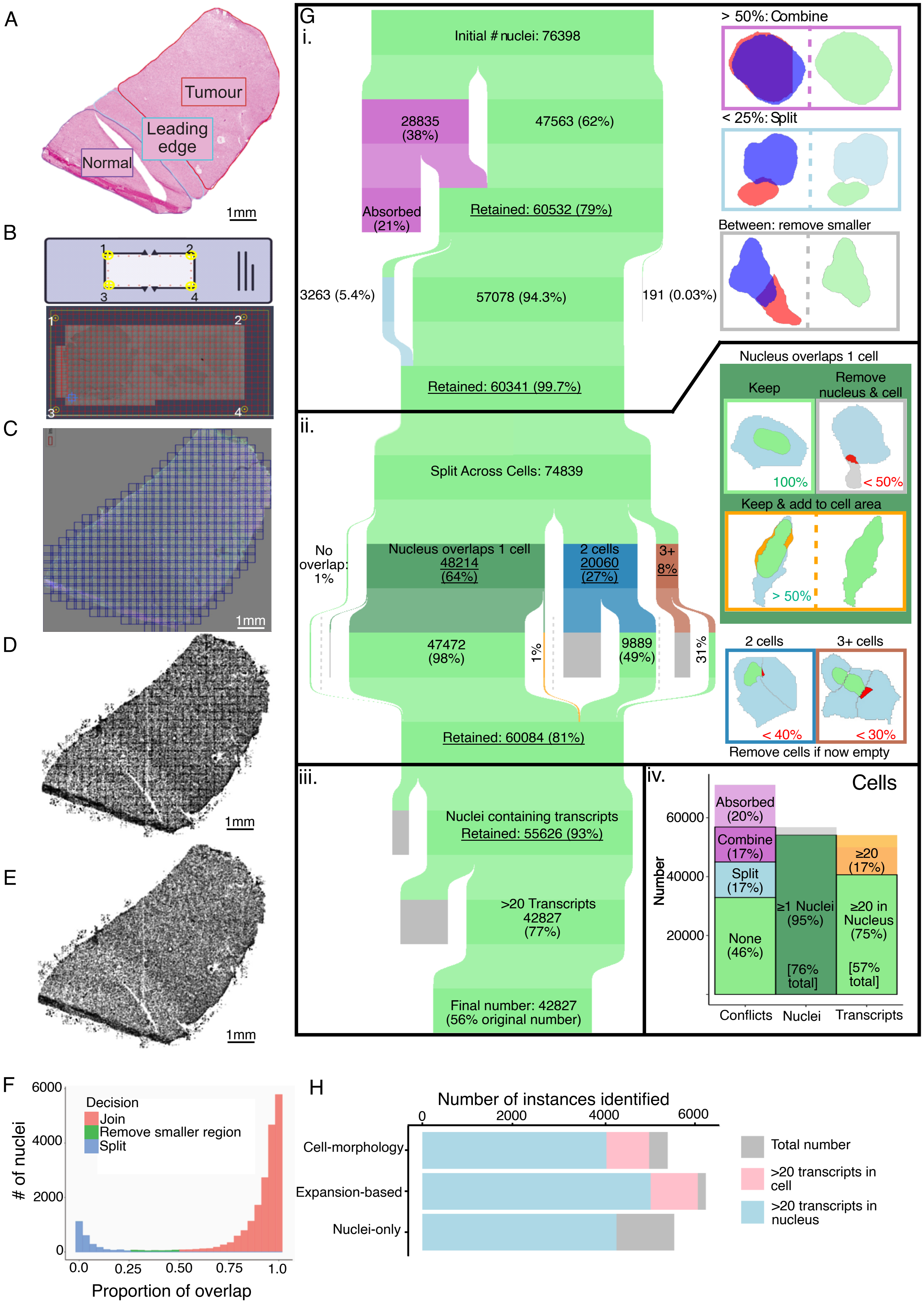

**Supplementary Figure 1.**

**A.** H&E of tumor tissue with annotation of histological regions. **B.** slide preparation protocol for imaging. **C.** Distribution of overlapping tiles used for segmentation. **D.** Localization of segmented nuclei before stitching. **E.** Localization of segmented nuclei after stitching. **F.** Histogram of proportion of overlap between overlapping nuclei, prior to stitching. Proportion is calculated as the percentage of area of the smaller nucleus that overlaps with the larger nucleus. Colored based on resulting stitching outcome based on overlap: join for >0.5, split for <0.25, and remove smaller for in-between. **G.** Sankey plot detailing the stepwise stitching protocol and its effect on final nuclei and cell numbers. i) Initial stitching protocol for nuclei-only segmentation, and amounts of nuclei affected by each decision, with examples for each shown on the right. ii) Consolidation of nuclei-only segmentations with cell-morphology segmentations, with examples for each decision point shown on the right. Iii) Transcript assignment to fully segmented nuclei and effects of subsequent filtration on final processed quantity. Iv) Proportions of cell-morphology segmentations affected by the various decision points. **H.** Comparison of number of segmentations identified with all three approaches: Cell-morphology, expansion-based, and nuclei-only segmentation.

Supplementary Figure 2

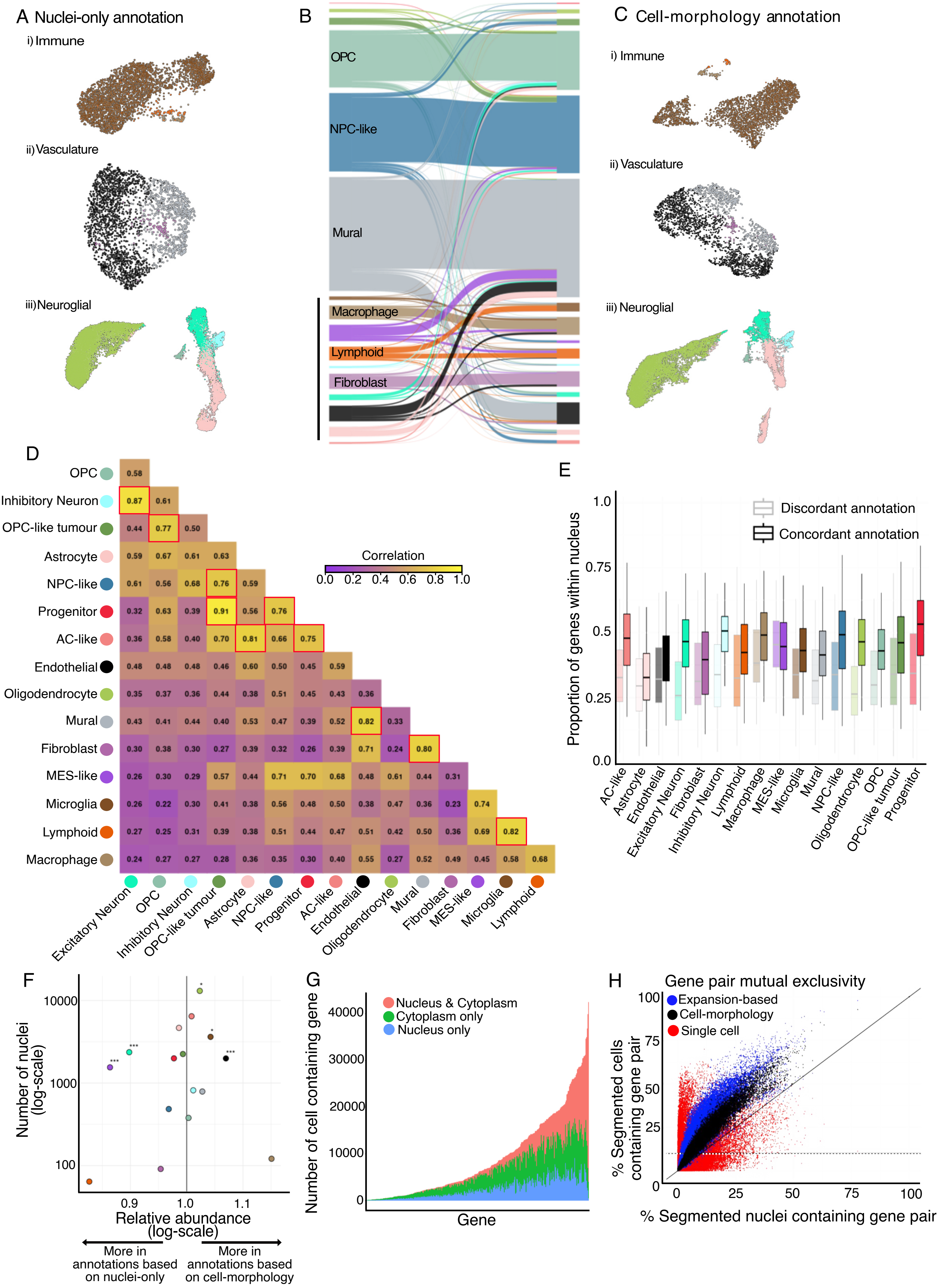

**Supplementary Figure 2.**

**A.** UMAP of nuclei-only segmentation annotated as i) immune cell types, ii) vasculature and iii) neuroglia, colored by nuclei-only annotation. **B.** Sankey plot showing change in annotation for matched nuclei-only segmentation (left) and cell-morphology segmentation (right) of lowly abundant cell types. Scale bar on bottom left, 1000 segmentations. **C.** UMAP of cell-morphology segmentation annotated as i) immune cell types, ii) vasculature and iii) neuroglia, colored by cell morphology annotation. **D.** Pairwise correlation of transcript expression profiles for each annotated cell type, based on nuclei-only segmentation. A higher correlation indicates a more similar cellular profile. Correlations above 0.75 are outlined in red. **E.** Proportion of transcripts bound to nucleus compared to cytoplasm, grouped and colored by nuclei-only cell type annotation, and split according to concordance with cell-morphology based annotation. **F.** Plot of relative abundance of cell types comparing annotations for nuclei-only and cell-morphology segmentation strategies (x-axis) compared to overall number of each cell type (y-axis, based on nuclei-only segmentation). P VALUE STATEMENT **G.** Assignment of genes to nucleus or cytoplasm. For each gene, every cell containing that gene is assessed for whether the gene is present in both the nucleus and cytoplasm, as determined by alternative expression profiles of nuclei-only and cell-morphology segmentation. Genes were ordered based on number of cells containing the gene form cell-morphology segmentation. **H.** Gene pair mutual exclusivity, calculated as the proportion of cells containing either gene that co-express both genes. X-axis, proportion identified with nuclei-only segmentation, Y-axis, proportion identified with various cell representations; either by cell-morphology (black), expansion-based (blue), or scRNA-seq (red). All gene pairs are shown, regardless of expression proportion in the single cell data.

Supplementary Figure 3

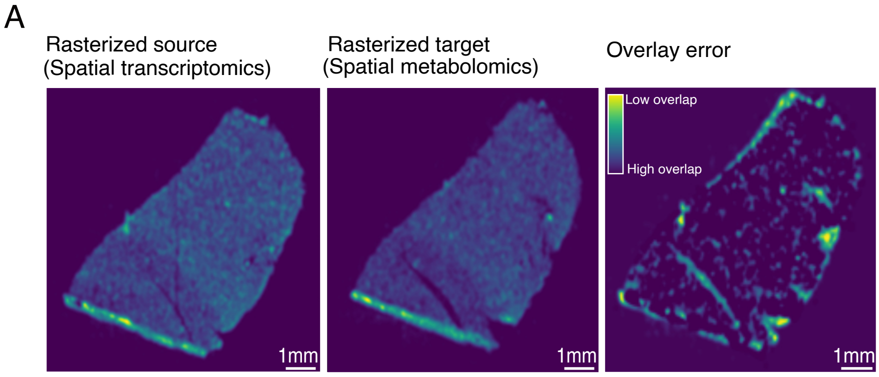


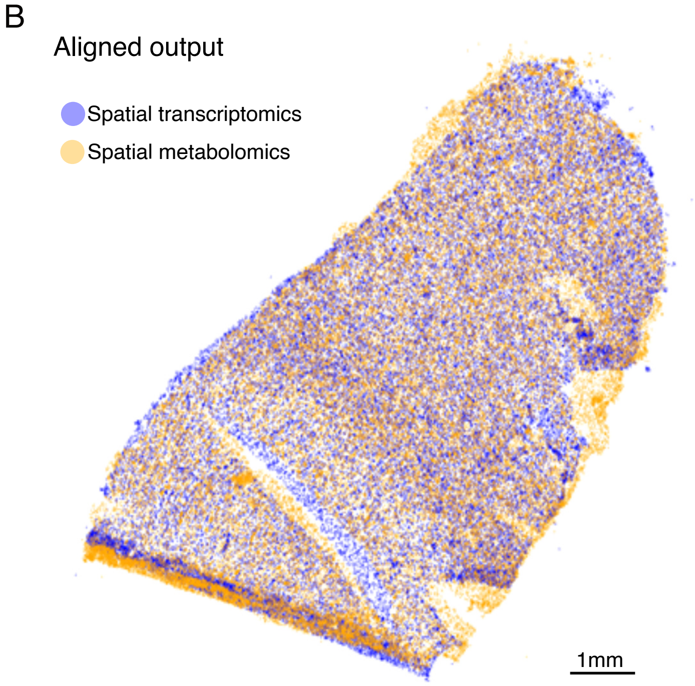

**Supplementary Figure 3.**

**A.** Rasterized points of source and target data prior to alignment including error map of alignment using STalign. Scale, 1mm. **B.** Aligned output of ST and SM data. Scale, 1mm.

Supplementary Figure 4

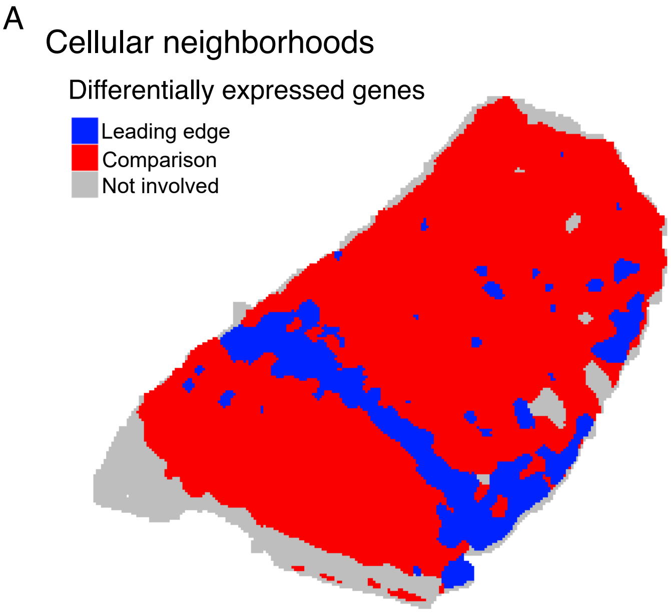


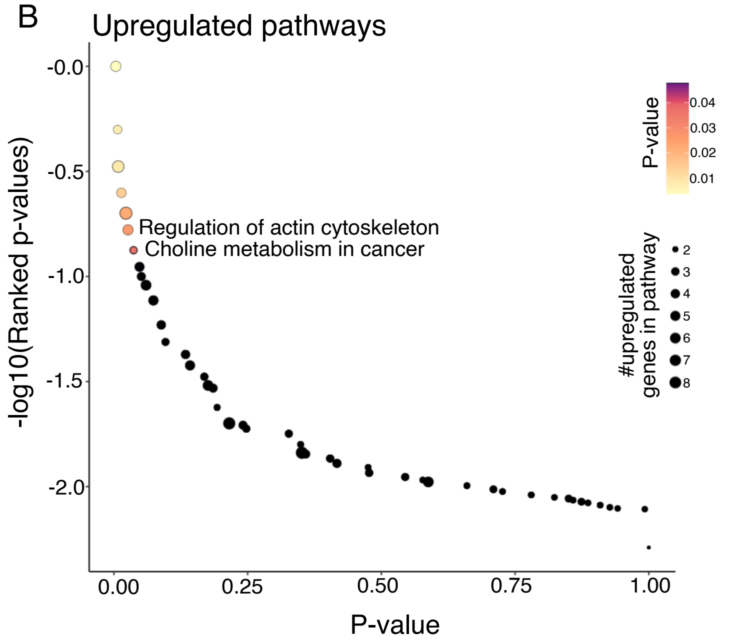


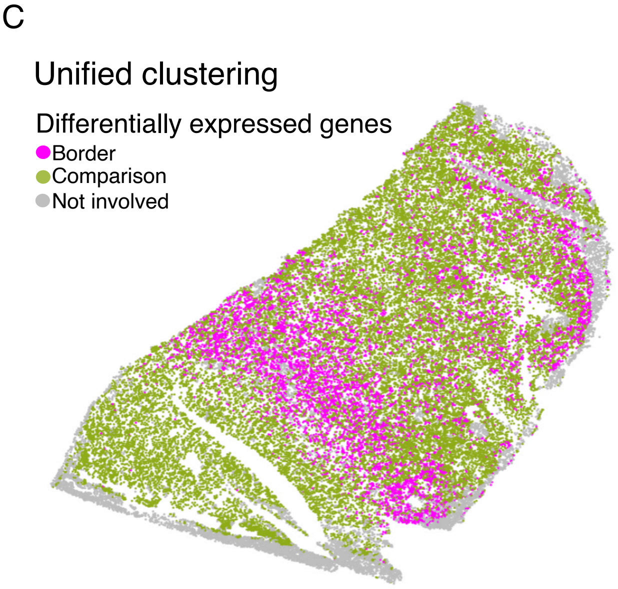


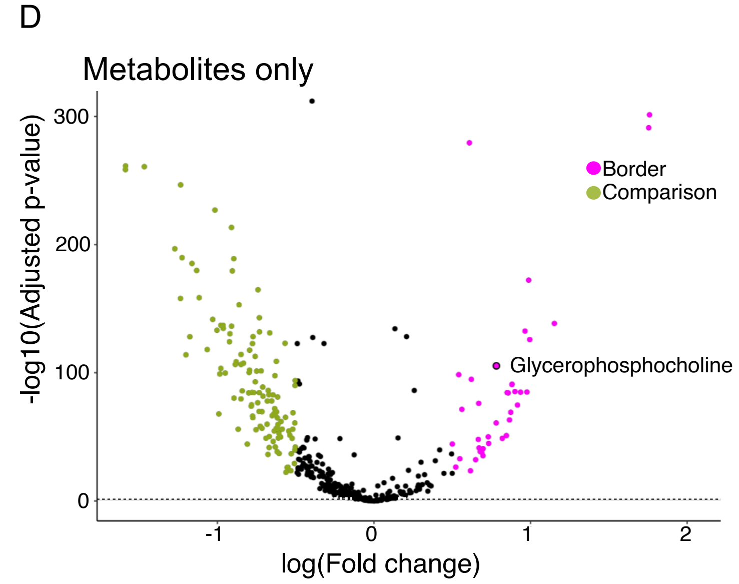

**Supplementary Figure 4.**

**A.** Spatial map of cellular neighborhoods colored by assignment for differential expression analysis of leading edge compared to normal and tumor core regions of tissue. **B.** Upregulated KEGG pathways in OPC-like tumor cells at the leading edge, compared to OPC-like genes in the normal or tumor core tissue regions. Ordered by p-value (x-axis) and overall rank of p-values (y-axis). **C.** Map of unified clustering of cells colored by assignment for differential expression analysis of leading edge compared to normal and tumor core regions of tissue. **D.** Volcano plot of differentially abundant metabolites at the leading edge, compared to the normal or tumor core tissue regions. Glycerophosphocholine (GPC) is highlighted.

**Supplementary Table 1.**

List of custom probes for spatial transcriptomics.
